# Kinetochore protein depletion underlies cytokinesis failure and somatic polyploidization in the moss *Physcomitrella patens*

**DOI:** 10.1101/438648

**Authors:** Elena Kozgunova, Momoko Nishina, Gohta Goshima

## Abstract

Lagging chromosome is a hallmark of aneuploidy arising from errors in the kinetochore– spindle attachment in animal cells. However, kinetochore components and cellular phenotypes associated with kinetochore dysfunction are much less explored in plants. Here, we carried out a comprehensive characterization of conserved kinetochore components in the moss *Physcomitrella patens* and uncovered a distinct scenario in plant cells regarding both the localization and cellular impact of the kinetochore proteins. Most surprisingly, knock-down of several kinetochore proteins led to polyploidy, not aneuploidy, through cytokinesis failure in >90% of the cells that exhibited lagging chromosomes for several minutes or longer. The resultant cells, containing two or more nuclei, proceeded to the next cell cycle and eventually developed into polyploid plants. As lagging chromosomes have been observed in various plant species in the wild, our observation raised a possibility that they could be one of the natural pathways to polyploidy in plants.

## Introduction

The kinetochore is a macromolecular complex that connects chromosomes to spindle microtubules and plays a central role in chromosome segregation. Kinetochore malfunction causes checkpoint-dependent mitotic arrest, apoptosis, and/or aneuploidy-inducing chromosome missegregation (1). Most of our knowledge on kinetochore function and impact on genome stability is derived from animal and yeast studies (2). Another major group of eukaryotes, plants, also possesses conserved kinetochore proteins (3–5). Although the localization and loss-of-function phenotype of some plant kinetochore proteins have been reported before (6–15), the data are mostly obtained from fixed cells of specific tissues. No comprehensive picture of plant kinetochore protein dynamics and functions can be drawn as of yet. For example, 12 out of 16 components that form CCAN (<u>c</u>onstitutive <u>c</u>entromere <u>a</u>ssociated <u>n</u>etwork) in animal and yeast cells cannot be identified by homology searches (2, 5). How the residual four putative CCAN subunits act in plants is also unknown.

The moss *Physcomitrella patens* is an emerging model system for plant cell biology. The majority of its tissues are in a haploid state, and, owing to an extremely high rate of homologous recombination, gene disruption and fluorescent protein tagging of endogenous genes are easy to obtain in the first generation (16). The homology search indicated that all the *P. patens* proteins identified as the homologue of human kinetochore components are conserved in the most popular model plant species *A. thaliana* (5): therefore, the knowledge gained in *P. patens* would be largely applicable to flowering plants, including crop species. Another remarkable feature of *P. patens* is its regeneration ability; for example, differentiated gametophore leaf cells, when excised, are efficiently reprogrammed to become stem cells (17, 18). Thus, genome alteration even in a somatic cell can potentially spread through the population.

In this study, we aimed to comprehensively characterize conserved kinetochore proteins in a single cell type, the *P. patens* caulonemal apical cell. We observed that many proteins displayed localization patterns distinct from their animal counterparts. Furthermore, kinetochore malfunction led to chromosome missegregation and microtubule disorganization in the phragmoplast, eventually resulting in cytokinesis failure and polyploidy.

## Results

### Endogenous localization analysis of conserved kinetochore proteins in *P. patens*

To observe the endogenous localization of putative kinetochore components, we inserted a fluorescent tag in-frame at the N- and/or C-terminus of eighteen selected proteins, which contain at least one subunit per sub-complex (Figure 1–Figure supplement 1). Initially we conducted C-terminal tagging since the success rate of homologous recombination is much higher than N-terminal tagging (19). For ten proteins, function was unlikely perturbed by tagging, as the transgenic moss grew indistinguishably from wild-type, despite the single-copy protein being replaced with the tagged protein. For other seven proteins, the functionality of the tagged version could not be verified, since untagged paralogs are present in the genome. The C-terminal tagging line for CENP-S could not be obtained after two attempts, suggesting that tagging affected the protein’s function and thereby moss viability. The N-termini of CENP-S, CENP-O, and CENP-C were also tagged with Citrine. Among them, no paralogous proteins could be identified for CENP-S or CENP-C; therefore, Citrine signals would precisely represent the endogenous localization. Exceptionally, histone H3-like CENP-A (CenH3) localization was determined by ectopic Citrine-CENP-A expression, as tagging likely perturbs its function.

**Figure 1.**
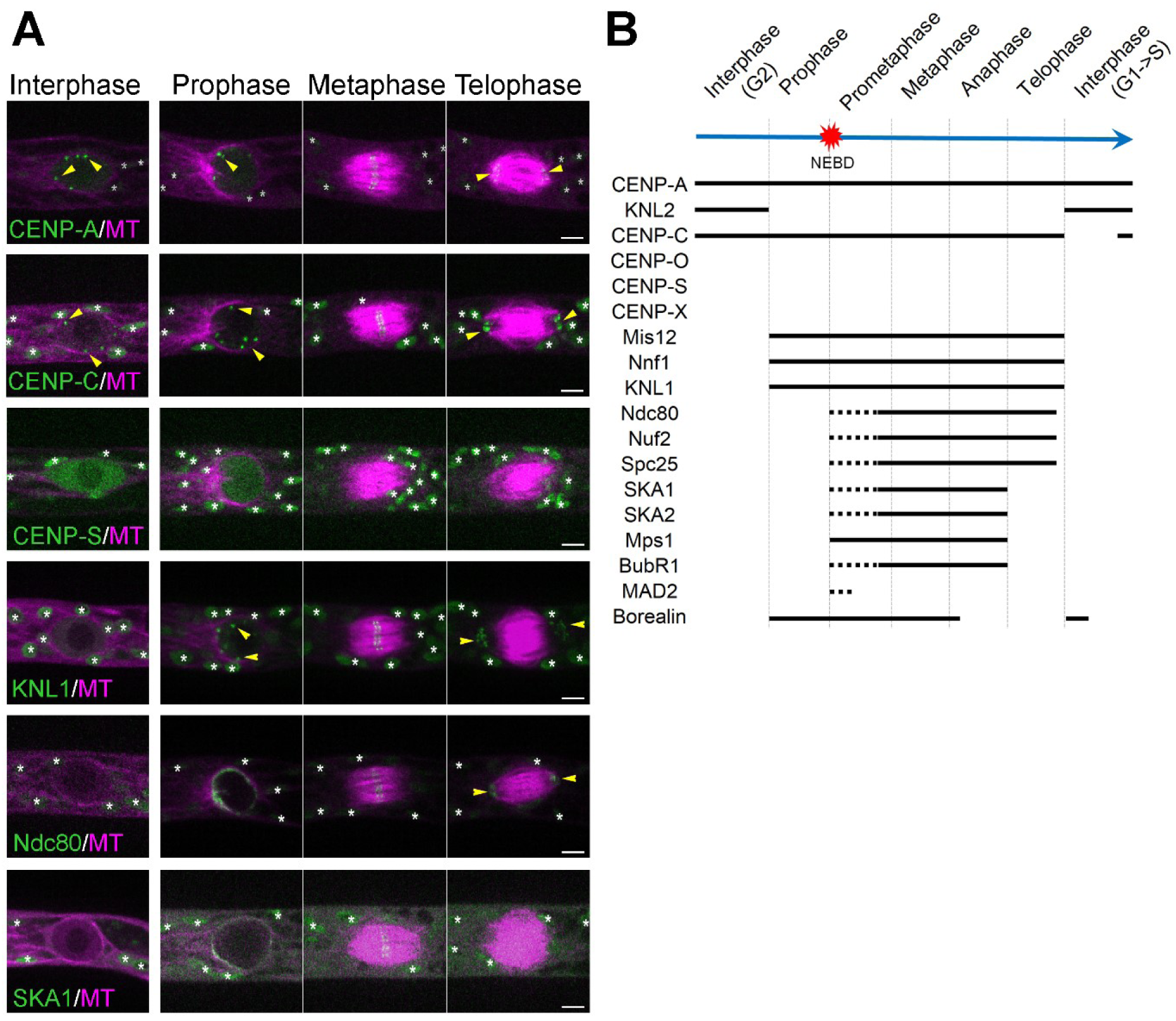
Unconventional localization of kinetochore proteins in *P. patens*. (A) Live imaging of *P. patens* caulonemal apical cells expressing mCherry-tubulin and selected kinetochore proteins: Citrine-CENP-A; Citrine-CENP-C; Citrine-CENP-S; KNL1-Citrine; Ndc80-Citrine and SKA1-Citrine. Full localization data can be found in Supplemental data. Some kinetochore signals are marked with yellow arrowheads, whereas autofluorescent chloroplasts are all marked with white asterisks. Images were acquired at a single focal plane. Bars, 5 µm. See Figure supplements 1–7, Video 1-4. (B) Timeline of kinetochore localization during the cell cycle in *P. patens* caulonemal apical cells. Solid lines correspond to the detection of clear kinetochore signals, whereas dotted lines indicate more dispersed signals.

Consistent with their sequence homology, many of the proteins were localized to the kinetochore at least transiently during the cell cycle. However, multiple proteins also showed unexpected localization (or disappearance) at certain cell cycle stages (Figure 1–Figure supplement 2–7; Video 1–4). Most surprising were CCAN protein dynamics: CENP-X, CENP-O and CENP-S did not show kinetochore enrichment at any stages (Figure 1–Figure supplement 3; Video 1, 3), whereas CENP-C also dissociated from the kinetochore transiently in the post-mitotic phase (Figure 1B–Figure supplement 4; Video 2, 3). Thus, we could not identify any “constitutive” kinetochore proteins other than CENP-A.

### Kinetochore malfunction causes chromosome missegregation and cytokinesis failure

We failed to obtain knockout lines and/or induce frameshift mutation using CRISPR/Cas9 for the single-copy kinetochore proteins, except for the spindle checkpoint protein Mad2, strongly suggesting that they are essential for moss viability. We therefore made conditional RNAi lines, targeting different proteins from both inner and outer kinetochores (summarized in Figure supplement 1). In this RNAi system, knockdown of target genes was induced by the addition of β-estradiol to the culture medium 4–6 days prior to live-imaging (20). Since RNAi sometimes exhibits an off-target effect, we prepared two independent RNAi constructs for most target genes. Following the previously established protocol (20, 21), we screened for cell growth/division phenotypes in ≥10 transgenic lines for each construct by using long-term (>10 h) fluorescent imaging. We observed mitotic defects in multiple RNAi lines, such as delay in mitotic progression, chromosome missegregation and/or multi-nuclei; these phenotypes were never observed in the control line (Figure 2A, B–Video 5). A full list of targeted genes and brief descriptions of the observed phenotypes are provided in Figure supplement 1.

**Figure 2.**
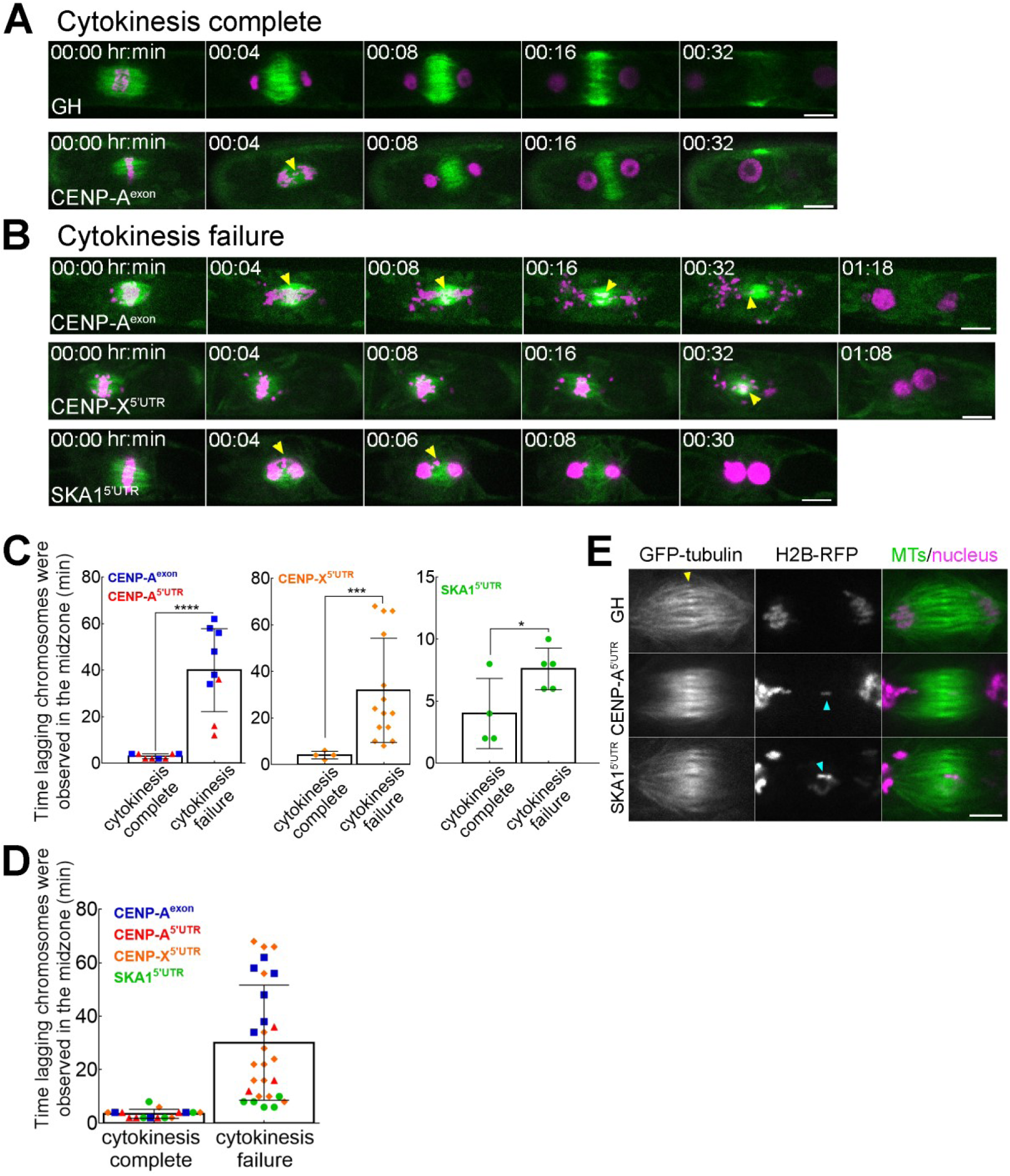
Lagging chromosomes in anaphase induce cytokinesis failure. (A, B) Lagging chromosomes (yellow arrowheads) present for several minutes in the midzone between separated chromatids cause cytokinesis failure in CENP-A, CENP-X and SKA1 RNAi lines. GH represents a control line. Bars, 10 µm. See Figure supplement 8–9, Video 5-8. (C, D) Correlation between cytokinesis failure and duration of lagging chromosomes observed in the midzone in the individual RNAi lines (C) and as combined data (D). Asterisks indicate significant differences between two groups (lagging chromosomes observed for short time or for several minutes) for two outcomes: cytokinesis complete and cytokinesis failure, calculated individually for CENP-A; CENP-X and SKA1 RNAi lines (*P = 0.0476, ***P = 0.0003, ****P < 0.0001; Fisher’s test; see Table supplement 1). Each data point corresponds to a single cell. Mean ± SD are presented. (E) Representative images of the microtubule overlap in the phragmoplast in the control line (GH) and in RNAi lines (CENP-A and SKA1) with lagging chromosomes. Note that microtubule overlaps appear more broad and fuzzy in RNAi cells. Yellow arrow indicates microtubule overlaps, whereas cyan arrows point to lagging chromosomes. Images were acquired with z-stacks and a single focal plane that best shows microtubule overlaps is presented. Bar, 5 µm.

We first selected CENP-A for detailed analysis, the only constitutive centromeric protein identified in *P. patens*. As expected, we observed a significant mitotic delay and chromosome alignment/segregation defects in the CENP-A RNAi lines (Figure 2–Figure supplement 8; Video 6). These phenotypes can be explained by a deficiency in proper kinetochore-microtubule attachment. Consequently, micronuclei were occasionally observed in the daughter cells, a hallmark of aneuploidy. We concluded that CENP-A, like in many organisms, is essential for equal chromosome segregation during mitosis in moss.

Surprisingly, we also frequently observed cells with two large nuclei in both RNAi lines (Figure 2B, 1 h 18 min), which is the typical outcome of cytokinesis failure in this cell type (22–24). To check if a similar phenotype is observed after the depletion of another kinetochore protein, we observed conditional RNAi line for SKA1, an outermost kinetochore component that does not directly interact with CENP-A and that had not been functionally characterized in the plant cells yet. As expected, mitotic delay and chromosome missegregation were observed in the RNAi line (Figure 2B–Figure supplement 8; Video 5). In addition, cytokinesis failure was also detected (Figure 2B–Video 7). To verify that the observed phenotype of SKA1 was not due to an off-target effect, we ectopically expressed RNAi-insensitive SKA1-Cerulean in the RNAi line and observed the rescue of all the above phenotypes (Figure 2–Figure supplement 9). Furthermore, we observed a similar phenotype in RNAi lines targeting CENP-C (CCAN), Nnf1 (Mis12 complex), KNL1 and Nuf2 (Ndc80 complex), suggesting that cytokinesis failure is a common outcome following kinetochore malfunction (Figure 2–Video 5).

Although we could not detect any kinetochore enrichment of the CCAN subunit CENP-X, we analyzed its RNAi lines. Interestingly, we observed similar phenotypes to CENP-A and SKA1, including cytokinesis failure (Figure 2B–Figure supplement 8; Video 6). CENP-X RNAi phenotypes were rescued by the ectopic expression of CENP-X-Cerulean that was resistant to the RNAi construct (Figure 2–Figure supplement 9). Thus, CENP-X has lost its kinetochore localization in moss, but is still essential for chromosome segregation and cell division.

By analyzing a total of 44 cells from SKA1 (9 cells), CENP-X (18 cells) and CENP-A RNAi (9 cells for one construct and 8 cells for the other) lines that had lagging chromosomes, we noticed a correlation between cytokinesis failure and lagging chromosomes lingering for a relatively long time in the space between separated chromatids. We therefore quantified the duration of lagging chromosomes’ residence in the midzone between separating chromatids following anaphase onset. Interestingly, a minor delay of chromosomes in the midzone (< 4 min) never perturbed cytokinesis (100%, n = 9 for CENP-A, n = 4 for CENP-X and n = 3 for SKA1). By contrast, if we observed a longer delay of chromosome clearance from the midzone, even when only a single chromosome was detectable, cytokinesis defects occurred in 96% of the cells (n = 9, 14 and 5; Figure 2C, D).

During plant cytokinesis, a bipolar microtubule-based structure known as the phragmoplast is assembled between segregating chromatids. The cell plate then forms in the phragmoplast midzone (∼4 min after anaphase onset in *P. patens* caulonemal cells) and gradually expands towards the cell cortex, guided by the phragmoplast (22). We observed that microtubules reorganized into phragmoplast-like structures upon chromosome segregation in every cell, regardless of the severity of chromosome missegregation (e.g. 32 min in Figure 2B). However, high-resolution imaging showed that microtubule interdigitates at the phragmoplast midzone were abnormal in the kinetochore RNAi lines. In 5 out of 7 control cells, a sharp microtubule overlap indicated by bright GFP-tubulin signals was observed during cytokinesis, as expected from previous studies (22, 25) (yellow arrowhead in Figure 2E). In contrast, CENP-A and SKA1 RNAi lines that had lagging chromosomes and eventually failed cytokinesis never exhibited such focused overlaps (0 out of 12 cells); instead, the overlap was broader and less distinguished (Figure 2E).

Finally, we checked if the cell plate was formed at any point in the cells that had cytokinesis defects, using the lipophilic FM4-64 dye. We could not observe vesicle fusion at the midzone following anaphase onset; thus, the cell plate did not form in the cells that had lagging chromosomes for a long time (Figure 2–Video 8). From these results, we concluded that occupation of the midzone by lagging chromosomes for several minutes prevents proper phragmoplast assembly and cell plate formation, which subsequently causes cytokinesis failure.

### Polyploid plants are regenerated from isolated multi-nucleated cells

Lagging chromosomes are a major cause of aneuploidy in daughter cells, which is particularly deleterious for haploid cells. However, the above observation supports a different scenario, whereby cytokinesis failure induced by lagging chromosomes allows a cell to have a duplicated genome set in two or more nuclei. On the other hand, whether animal somatic cells that have failed cytokinesis can re-enter the cell cycle or not remains an ongoing debate (26–28). To address whether moss cells can recover from severe cell division defects and continue their cell cycle, we first analyzed the DNA content of cells in the CENP-A exon-targeting RNAi line, in which multi-nucleated cells were most prevalent. For comparison, we used the parental line: the nuclei of anaphase/telophase cells served as the 1N reference and randomly selected interphase nuclei as the 2N reference, as caulonemal cells are mostly in the G2 phase (18, 29). We observed that the majority of the multi-nucleated cells after CENP-A RNAi underwent DNA replication and became tetraploid or attained even higher ploidy (Figure 3A; DNA was quantified at day 5 after β-estradiol treatment).

**Figure 3.**
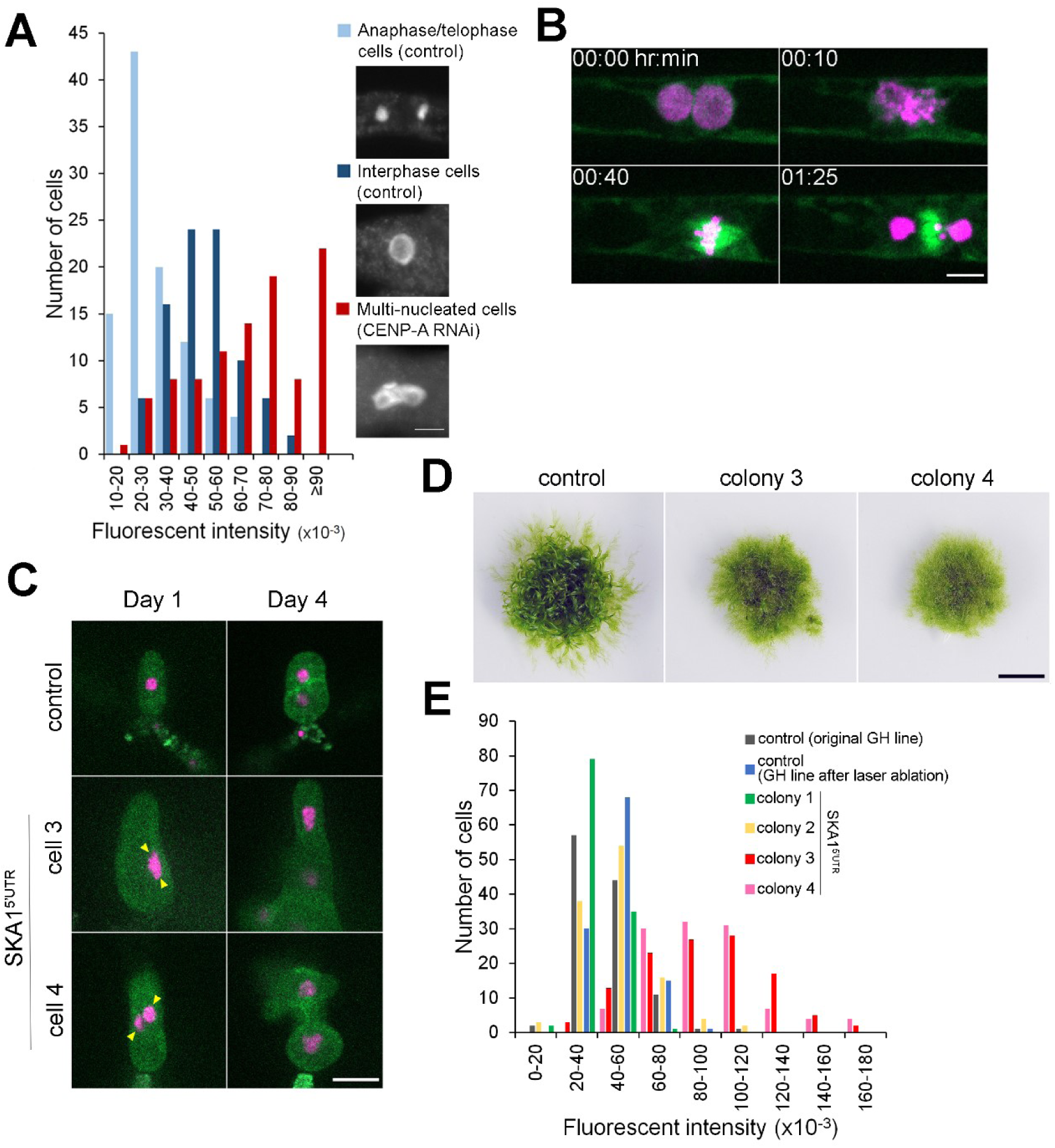
Cytokinesis failure in somatic cells can generate plants with whole-genome duplication. (A) Quantification of the nuclear DNA content. Anaphase/telophase cells were used as a standard for 1N nuclei (light blue). Interphase cells randomly selected in the control line mostly had double amounts of DNAs as expected (dark blue), whereas cells that failed cytokinesis had higher ploidy (red). DNA amounts are shown as fluorescent intensity of the DAPI-stained nuclei per cell after subtraction of the cytoplasmic background. (B) Representative images of mitotic entry and single spindle formation of the multi-nucleated cell in the *P. patens* SKA1 RNAi line. Bar, 5 µm. See Video 9. (C) Regeneration of a single cell isolated by laser dissection microscopy from the control cell line (GH) or multi-nucleated cells from SKA1 RNAi line (multi-nuclei are marked with yellow arrowheads). Bar, 50 µm. (D) Moss colonies regenerated from single cells. Bar, 0.5 cm. (E) Quantification of the nuclear DNA content in the interphase nucleus of regenerated moss colonies, corresponding to (C) and (D).

Next, we checked if multi-nucleated cells continue cell cycling. We used SKA1 RNAi line for a long (46 h) time-lapse imaging; with this imaging, we expected to monitor the process of cytokinesis failure of a haploid cell and its fate. During the imaging period, we indeed observed cytokinesis failure and 10% or 25% multi-nucleated apical cells executed the next cell division by forming a single spindle (n = 43 and 25 for experiments 1 and 2, respectively, Figure 3B; Video 9). The reason for the low frequency of this event is unclear; strong chromosome missegregation might result in a severe “aneuploid” state for each nucleus, whereas the cell is overall polyploid, which might change the cell physiology. Nevertheless, this data strongly suggests that cells that have undergone cytokinesis failure can continue cell cycling as diploids at a certain probability.

Diploid *P. patens* is known to develop protonema tissue with a few gametophores (leafy shoots) (30); therefore, a multi-nucleated cell produced by the cytokinesis failure of a caulonemal cell might proliferate and form a large protonema colony. To test this possibility, we isolated and cultured several cells (Figure 3C) that were seemingly multi-nuclear after SKA1 RNAi via laser dissection microscopy (note that there is an unambiguity in identifying multi-nucleate cells; see Methods for detailed explanation). After 6 weeks of culturing without β-estradiol (i.e. RNAi was turned off), we obtained four moss colonies, two of which consisted mainly of protonemal cells with a few gametophores (Figure 3D, colony 3 and 4). DNA staining and quantification showed that the majority of the cells derived from those two colonies had DNA content approximately double of the control haploid cells, which were regenerated in an identical manner (Figure 3E, colony 3 and 4, regenerated from cell 3 and 4, respectively). Thus, a polyploid plant was regenerated from a single multi-nucleated somatic cell.

## Discussion

### Kinetochore protein dynamics in a plant cell

This study provides a comprehensive view of the dynamics of conserved kinetochore proteins in a single cell type of *P. patens*; furthermore, to the best of our knowledge, several proteins, including borealin, KNL1 and SKA subunits, have been characterized for the first time in plant cells. The tagged proteins were expressed under their native promoter at the original chromosome locus; thus, fluorescent signals of most, if not all, proteins would represent the endogenous localization.

Overall, the behavior of outer subunits was largely consistent with their animal counterparts, suggesting that the mitotic function is also conserved. However, the timing of kinetochore enrichment differed from that of animal cells and even flowering plants (e.g. Arabidopsis, maize) (6, 14, 31): for example, *P. patens* Ndc80 complex gradually accumulated at the kinetochore after NEBD, unlike Arabidopsis and maize, where it showed kinetochore enrichment throughout the cell cycle (6, 14). More unexpected localizations were observed for inner CCAN subunits, namely CENP-C, CENP-O, CENP-S and CENP-X. For example, CENP-C disappeared from the centromeres shortly after mitotic exit. In animal cells, CENP-C has been suggested to act in cooperation with Mis18BP1/KNL2 to facilitate CENP-A deposition in late telophase and early G1 (2). Hence, the mechanism of CENP-A incorporation might have been modified in plants.

CENP-O, -S, or –X did not show kinetochore enrichment at any stage. CENP-X localization was unlikely an artifact of Citrine tagging, since the tagged protein rescued the RNAi phenotype. In human cells, sixteen CCAN subunits, forming four sub-complexes, have been identified and shown to be critical for kinetochore assembly and function, not only in cells, but also in reconstitution systems (32, 33). In plants, only four CCAN homologues have been identified through sequence homology search. It is therefore possible that less conserved CCAN subunits are present, but could not be identified by the homology search. However, the complete lack of kinetochore localization for CENP-O, -S, -X suggests that plants have lost the entire kinetochore-enriched CCAN complex. Somewhat puzzlingly, CENP-X, despite its unusual localization, remained an essential factor for chromosome segregation in *P. patens*. In animals, it has been proposed that CENP-S and CENP-X form a complex and play an important in outer kinetochore assembly (34). It is an interesting target for further investigation if plant CENP-S/CENP-X preserves such a function.

### Chromosome missegregation causes polyploidization

We observed lagging chromosomes as well as cytokinesis failure after knocking down kinetochore components. Failure in chromosome separation/segregation and cytokinesis can be caused by a single gene mutation, if the gene has multiple functions; for example, separase Rsw4 (*radially swollen4*) in *A. thaliana* is involved in sister chromatid separation, cyclin B turnover and vesicle trafficking that is required for phragmoplast formation (35–38). By contrast, in our study, both phenotypes were observed after RNAi treatment of CENP-A, a constitutive centromeric histone protein that is unlikely to play a direct role in cytokinesis. Furthermore, the cytokinesis phenotype frequently appeared in RNAi lines targeting other six kinetochore proteins, and only when lagging chromosomes were present. Based on these data, we propose that persistent lagging chromosomes cause cytokinesis failure. Lagging chromosomes might act as physical obstacles to perturb phragmoplast microtubule amplification and/or cell plate formation. Alternatively, persistent lagging chromosomes might produce an unknown signal or induce a certain cell state that inhibits phragmoplast expansion and/or cell plate formation in order to prevent chromosome damage, reminiscent of the NoCut pathway in animal cytokinesis (39, 40). We favor the latter model, as abnormal microtubule interdigitates were observed in the whole phragmoplast and not limited to the region proximal to the lagging chromosome (Figure 2E). Notably, in a recent study, cytokinesis in moss protonema cells could be completed despite longer microtubule overlaps (41). It suggests that abnormal microtubule interdigitates represent the consequence of microtubule dynamics mis-regulation rather than the direct cause of cytokinesis failure.

Our data further suggest that, in *P. patens*, chromosome missegregation in a single cell could lead to the generation of polyploid plants. Could lagging chromosomes cause polyploidization through somatic cell lineage in wild-type plants? In our imaging of control moss cells, we could not find any lagging chromosome, since mitotic fidelity is very high in our culture conditions. Intriguingly, however, various mitotic abnormalities, including lagging chromosomes have been long observed in wild-type plants and crops, albeit at a low frequency and/or under harsh natural conditions (42–44). Those studies did not analyze the relationship between lagging chromosomes and cytokinesis integrity; we expect the presence of lagging chromosomes for a certain duration to similarly perturb cytokinesis as observed in our study of moss, since the cytokinesis process is highly conserved between bryophytes and angiosperms (45). Genome sequencing suggests that *P. patens*, like many other plant species, experienced whole genome duplication at least once during evolution (46). Polyploidization through spontaneous mitotic errors in somatic cells might have a greater impact on *de novo* formation of polyploid plants than previously anticipated.

## Materials and Methods

### Moss culture and transformation

We generally followed protocols described by Yamada *et al* (19). In brief, *Physcomitrella patens* culture was maintained on BCDAT medium at 25°C under continuous light. Transformation was performed with the polyethylene glycol-mediated method and successful endogenous tagging of the selected genes was confirmed by PCR (19). We used *P. patens* expressing mCherry-α-tubulin under the pEF1α promoter as a host line, except for Mis12-mCherry line where GFP-α-tubulin line was used as a host line. For knockout, CRISPR (47) and RNAi transformations, we used the GH line, expressing <u>G</u>FP-tubulin and <u>H</u>istoneH2B-mRFP. *P. patens* lines developed for this study are described in Dataset S1.

### Plasmid construction

Plasmids and primers used in this study are listed in Dataset S2. For the C-terminal tagging, we constructed integration plasmids, in which ∼800 bp C-terminus and ∼800 bp 3’-UTR sequences of the kinetochore gene were flanking the *citrine* gene, the nopaline synthase polyadenylation signal (nos-ter), and the G418 resistance cassette. For the N-terminal tagging we constructed integration plasmids, in which ∼800 bp 5’-UTR and ∼800 bp N-terminus sequences of the kinetochore gene were flanking the *citrine* gene. CENP-A cDNA was amplified by PCR and sub-cloned into a vector containing the rice actin promoter, *citrine* gene, the rbcS terminator, the modified *aph4* cassette, and flanked by the genomic fragment of the *hb7* locus to facilitate integration. All plasmids were assembled with the In-Fusion enzyme according to manufacturer’s protocol (Clontech). RNAi constructs were made by using the Gateway system (Invitrogen) with pGG624 as the destination vector (21).

### DNA staining

We followed the protocol described by Vidali *et al* (48) with the following modifications: sonicated moss was cultured for 6–7 days on the BCDAT plate, containing 5 µM β-estradiol for RNAi induction and 20 µg/ml G418 to prevent contamination. Collected cells were preserved in a fixative solution (2% formaldehyde, 25 mM PIPES, pH 6.8, 5 mM MgCl_2_, 1 mM CaCl_2_) for 30 min and washed three times with PME buffer (25 mM PIPES, pH 6.8, 5 mM MgCl_2_, 5 mM EGTA). Following fixation, cells were mounted on 0.1% PEI (polyethyleneimine)-coated glass slides and subsequently incubated with 0.1% Triton X-100 in PME for 30 min and 0.2% driselase (Sigma-Aldrich) in PME for 30 min. Next, cells were washed twice in PME, twice in TBS-T buffer (125 mM NaCl, 25 mM Tris-HCl, pH 8, and 0.05% Tween 20) and mounted in 10 µg/mL DAPI in TBS-T for observation. Images were acquired with the Olympus BX-51 fluorescence microscope equipped with ZEISS Axiocam 506 Color and controlled by ZEN software. Fluorescent intensity was measured with ImageJ. Cytoplasmic background was subtracted.

### Live-imaging microscopy

A glass-bottom dish (Mattek) inoculated with moss was prepared as described in Yamada et al (19) and incubated at 25°C under continuous light for 4–7 days before live-imaging. To observe RNAi lines, we added 5 µM β-estradiol to culture medium (21). For the high magnification time-lapse microscopy, the Nikon Ti microscope (60×1.40-NA lens or 100×1.45-NA lens) equipped with the spinning-disk confocal unit CSU-X1 (Yokogawa) and an electron-multiplying charge-coupled device camera (ImagEM; Hamamatsu) was used. Images were acquired every 30 s for localization analysis and every 2 min for RNAi analysis. The microscope was controlled by the Micro-Manager software and the data was analyzed with ImageJ. The rescue lines for RNAi were observed using a fluorescence microscope (IX-83; Olympus) equipped with a Nipkow disk confocal unit (CSU-W1; Yokogawa Electric) controlled by Metamorph software.

### Single cell isolation

Protonema tissue of *P. patens* was sonicated, diluted with BCD medium with 0.8% agar, and spread on cellophane-covered BCDAT plates that contain 5 µM estradiol to induce RNAi. After 5–6 days, small pieces of cellophane containing clusters of protonemal cells (each containing 3–20 cells) were cut with scissors and placed upside-down on a glass-bottom dish. Bi- or multi-nucleated cells were identified using Axio Zoom.v16. Single bi-nucleated cell (SKA1 RNAi line) or random cell (control GH line) was selected for isolation and all other cells were ablated with a solid-state ultraviolet laser (355 nm) through a 20X objective lens (LD Plan-NEOFLUAR, NA 0.40; Zeiss) at a laser focus diameter of less than 1 µm using the laser pressure catapulting function of the PALM microdissection system (Zeiss). Irradiation was targeted to a position distantly located from the cell selected for isolation to minimize the irradiation effect. Note that visual distinction of multi-nucleated cells from those with slightly deformed nuclei is not easy in *P. patens*, since in multi-nucleated cells, the nuclei maintain very close association with each other, so that nuclear boundaries often overlap. We interpret that two of four regenerated protonemata had haploid DNA content due to our unintentional isolation of a single cell with a deformed nucleus rather than multi-nuclei. Next, a piece of cellophane with single isolated cell was transferred from the glass-bottom dish to estradiol-free medium (20 µg/ml G418 was supplied to prevent bacterial/fungal contamination). DAPI staining was performed 5–6 weeks later as described above.

### Sequence analysis

Full-size amino acid sequences of the selected proteins were aligned using MAFFT ver. 7.043 and then revised manually with MacClade ver. 4.08 OSX. We used the Jones-Taylor-Thornton (JTT) model to construct maximum-likelihood trees in MEGA5 software. Statistical support for internal branches by bootstrap analyses was calculated using 1,000 replications. Reference numbers correspond to Phytozome (www.phytozome.net) for *Physcomitrella patens*, the Arabidopsis Information Resource (www.arabidopsis.org) for *Arabidopsis thaliana* and Uniprot (www.uniprot.org) for *Homo sapiens*. Original protein alignments after MAFFT formatted with BoxShade (https://embnet.vital-it.ch/software/BOX_form.html) are shown in Supplemental dataset 3.

## Acknowledgements

We are grateful to Dr. Yoshikatsu Sato and Nagisa Sugimoto for their assistance with laser ablation experiments; to Dr. Peishan Yi, Moé Yamada and Shu Yao Leong for comments and discussion; and Rie Inaba for technical assistance. Imaging was partly conducted in the Institute of Transformative Bio-Molecules (WPI-ITbM) at Nagoya University, supported by Japan Advanced Plant Science Network. This work was funded by JSPS KAKENHI (17H06471, 17H01431) to G.G. The authors declare no competing financial interests.

## Supplemental materials

### Abbreviations

CCAN: <u>C</u>onstitutive <u>C</u>entromere <u>A</u>ssociated <u>N</u>etwork
Cit: Citrine
CPC: <u>C</u>hromosome <u>P</u>assenger <u>C</u>omplex
GFP: <u>g</u>reen fluorescent <u>p</u>rotein
HR: <u>h</u>omologous recombination
mCh: mCherry
MTs: microtubules
NEBD: <u>n</u>uclear <u>e</u>nvelope <u>b</u>reak<u>d</u>own
RNAi: RNA interference
SAC: <u>s</u>pindle <u>a</u>ssembly <u>c</u>heckpoint

## Full names

BMF1 *(Bub1)* - <u>B</u>UB1/<u>M</u>AD3 family 1

BubR1 (BMF2) - <u>BUB</u>1-related protein <u>1</u> (BUB1/MAD3 family 2)

CENP-A *(cenH3)* – <u>cen</u>tromere <u>p</u>rotein A *(centromeric Histone 3)*

CENP-C – <u>cen</u>tromere <u>p</u>rotein C

CENP-O – <u>cen</u>tromere <u>p</u>rotein O

CENP-S *(FAAP16, MHF1)* – Centromere protein S *(Fanconi anemia-associated polypeptide of 16 kDa; FANCM-associated histone fold protein 1)*

CENP-X *(FAAP10, MHF2)* – Centromere protein X *(Fanconi anemia-associated polypeptide of 10 kDa; FANCM-associated histone fold protein 2)*

Dsn1 – <u>d</u>osage <u>s</u>uppressor of <u>N</u>nf<u>1</u>

KNL1 *(Spc7; Blinkin)* – <u>k</u>inetochore <u>n</u>ull <u>1</u> *(spindle pole body component 7; Bub-linking kinetochore protein)*

KNL2 *(MIS18BP1)* – <u>k</u>inetochore <u>n</u>ull <u>2</u> *(Mis18-binding protein 1)*

MAD2 - <u>m</u>itotic <u>a</u>rrest <u>d</u>eficient <u>2</u>

Mis12 – <u>m</u>inichromosome in<u>s</u>tability <u>12</u>

Mps1 – serine/threonine-protein kinase MPS1 (<u>m</u>onopolar <u>s</u>pindle <u>p</u>rotein <u>1</u>)

Ndc80 *(HEC1)* - <u>n</u>uclear <u>d</u>ivision <u>c</u>ycle protein <u>80</u> *(highly expressed in cancer 1)*

Nnf1*(PMF1)* - <u>n</u>ecessary for <u>n</u>uclear function <u>1</u> *(Polyamine-modulated factor 1)*

Nuf2 - <u>nu</u>clear filament-containing protein <u>2</u>

SKA1, 2, 3 – <u>s</u>pindle and <u>k</u>inetochore <u>a</u>ssociated protein 1, 2, 3

Spc24, 25 - <u>s</u>pindle <u>p</u>ole body <u>c</u>omponent 24, 25

Taf9 - <u>TA</u>TA box binding protein (TBP)-associated factor

---

**Figure supplement 1.**
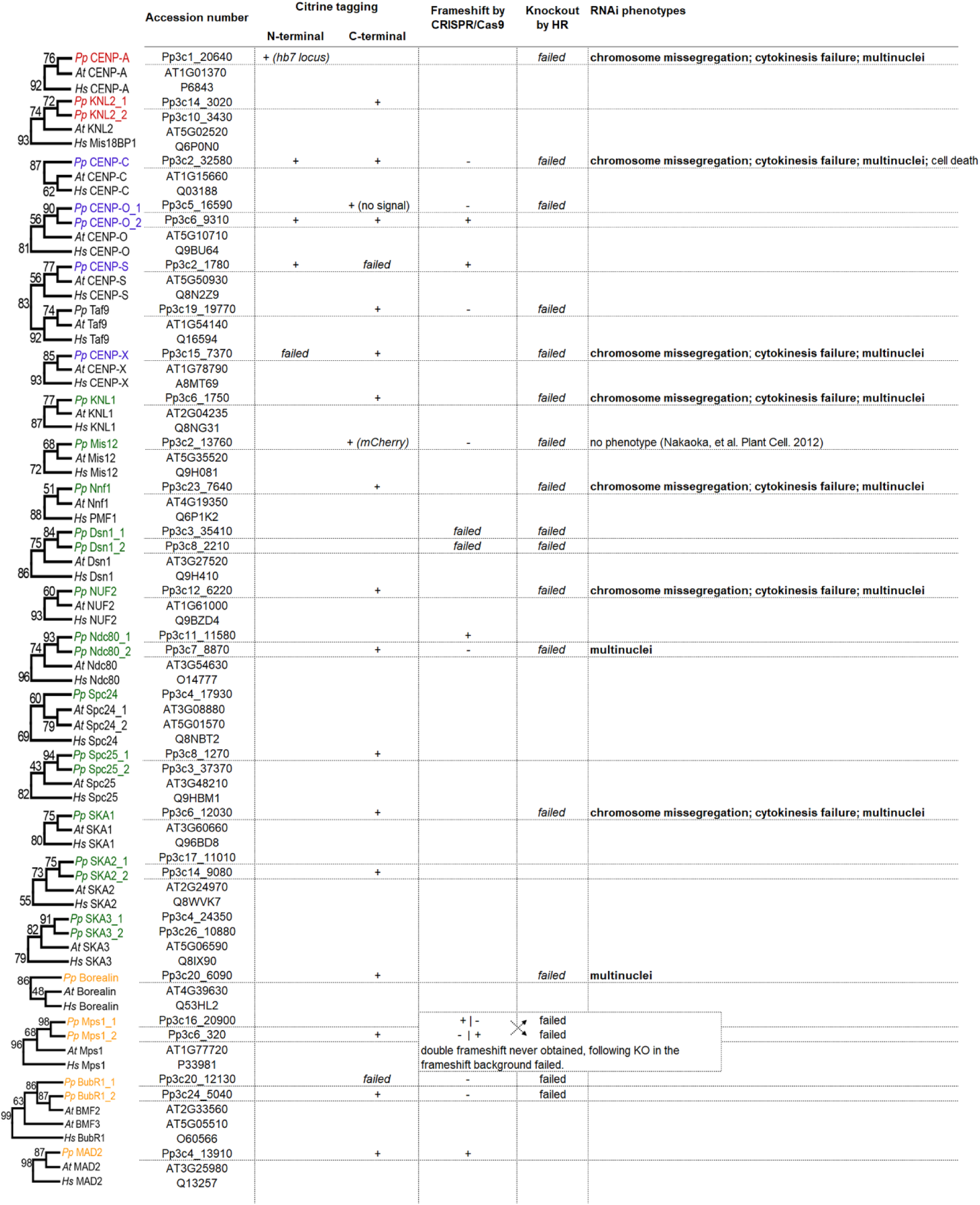
Summary of kinetochore protein tagging and disruption/knockdown in *P. patens*. *(Left)* Maximum-likelihood phylogenetic trees of conserved centromere/kinetochore proteins in *Physcomitrella patens*, *Arabidopsis thaliana* and *Homo sapiens*. Numbers represent bootstrapping values (above 50%) calculated from 1,000 replications. Accession numbers for each protein correspond to Phytozome (https://phytozome.jgi.doe.gov/) for *P. patens*; TAIR (https://www.arabidopsis.org/) for *A. thaliana* and UniProt (http://www.uniprot.org/) for *H. sapiens*. *(Middle)* Summary of Citrine tagging pursued in this study. (*Right*) Summary of knockout, CRISPR/Cas9 frameshift (“-” indicates that frameshift mutations could not be obtained) and RNAi experiments pursued in this study. HR stands for homologous recombination. “+” indicates successful transgenic line selection.

**Figure supplement 2.**
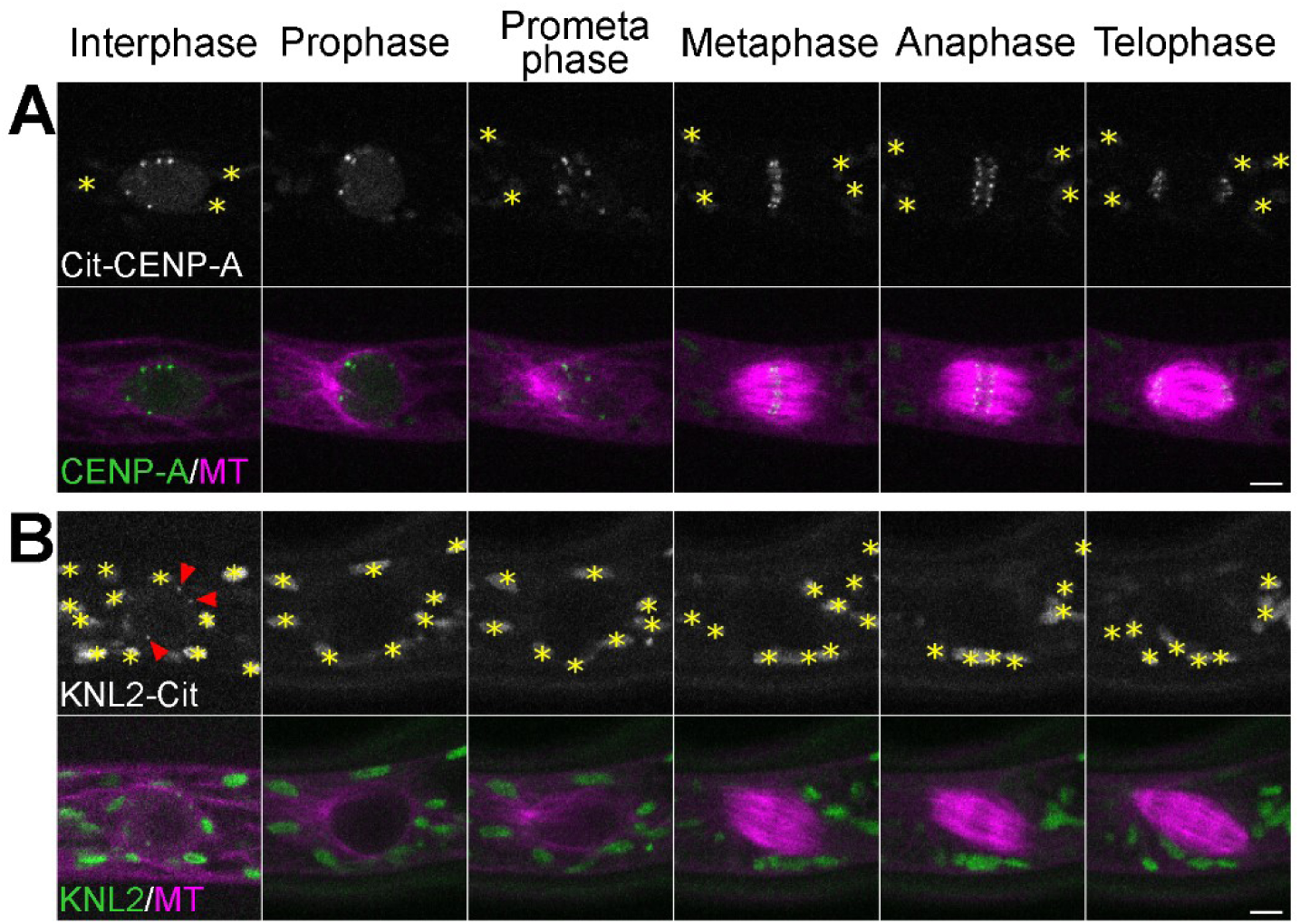
Localization of CENP-A and KNL2/MIS18BP1 during cell division. Live imaging of *P. patens* protonemal apical cells expressing mCherry-tubulin (magenta) and Citrine-CENP-A (A) or KNL2-Citrine (B). Citrine-CENP-A data is an expanded version of Figure 1. Autofluorescent chloroplasts are marked with yellow asterisks. Images were obtained at a single focal plane. CENP-A was localized at the centromeric region throughout the cell cycle, whereas KNL2-Citrine was visible only during interphase (red arrowheads). Bars, 5 µm.

**Figure supplement 3.**
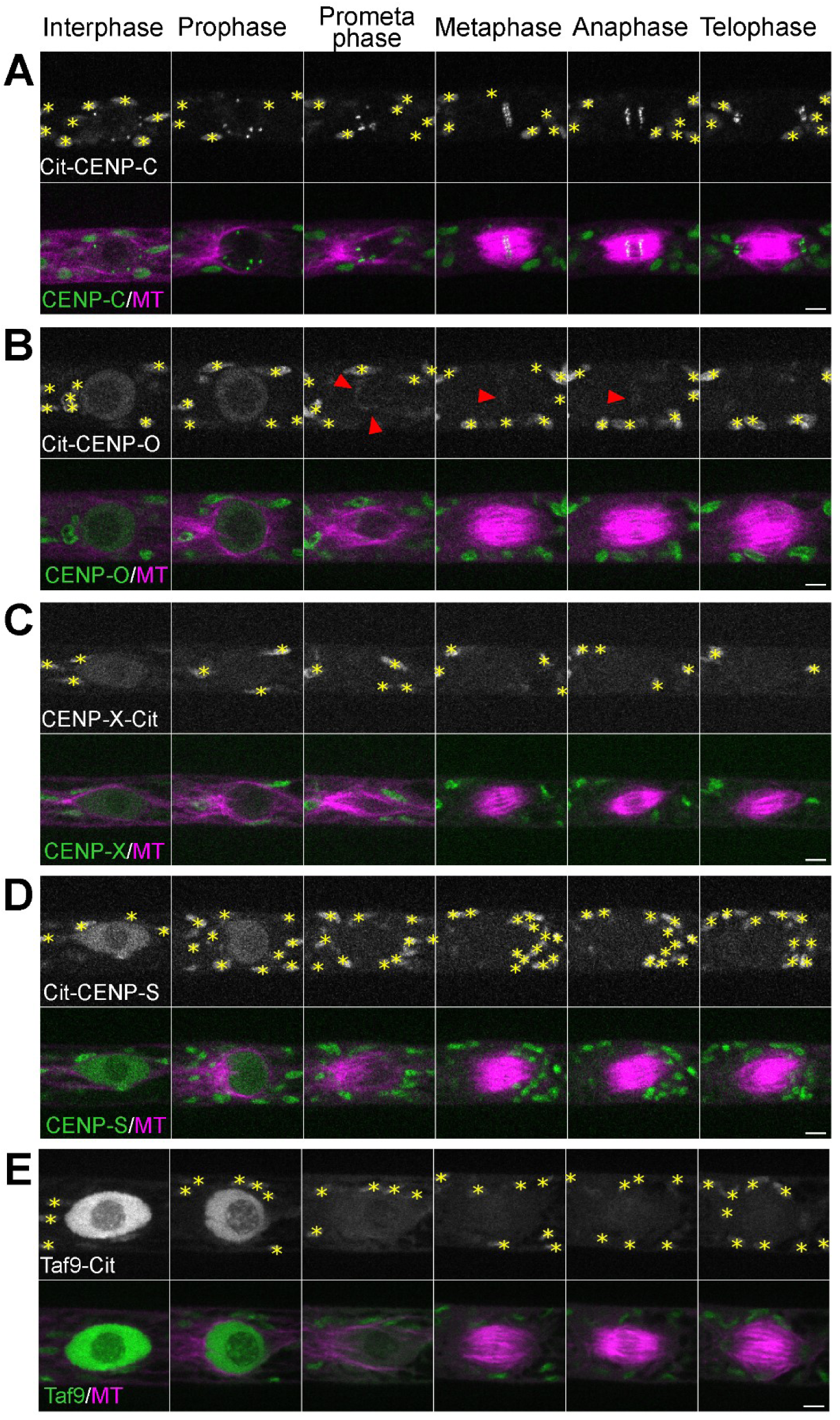
Localization of CCAN proteins during cell division. Live imaging of *P. patens* protonemal apical cells expressing mCherry-tubulin (magenta) and Citrine-tagged (green) CENP-C (A), CENP-O (B), CENP-X (C), CENP-S (D) and CENP-S-like protein Taf9 (E). Citrine-CENP-C and Citrine-CENP-S data are expanded versions of Figure 1. Autofluorescent chloroplasts are marked with yellow asterisks. Images were obtained at a single focal plane. CENP-C was localized at the centromere from G2 to telophase, whereas none of the other CCAN proteins showed punctate signals throughout the cell cycle. CENP-O showed weak midzone localization from prometaphase to anaphase (arrowheads). Bars, 5 µm.

**Figure supplement 4.**
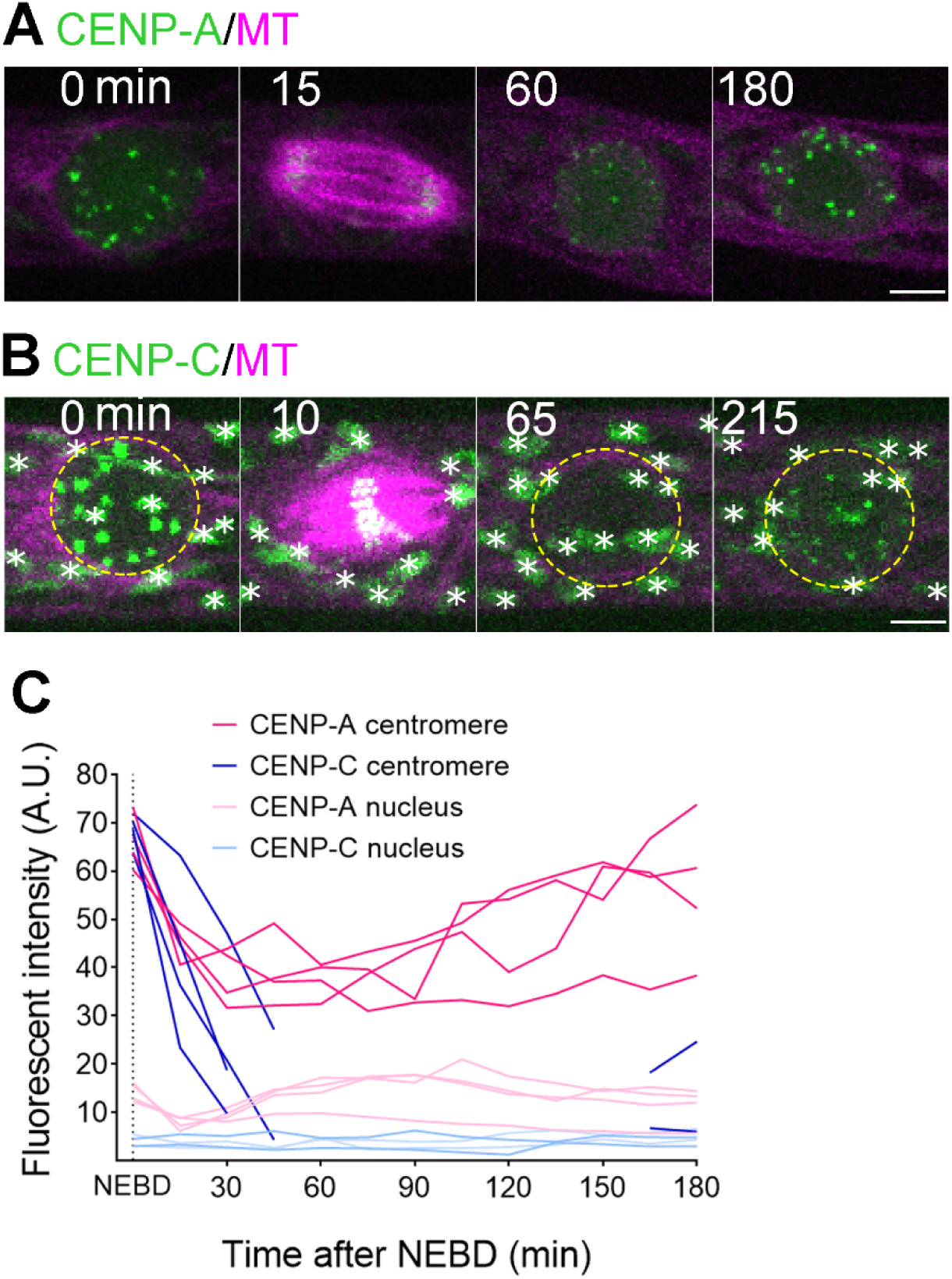
CENP-C is not a constitutive centromeric protein in *P. patens*. Citrine-CENP-A (A) and Citrine-CENP-C (B) localization starting from NEBD. At each time point, ten z-sections were acquired (separated by 1 µm). Merged images of mCherry-tubulin (single focal plane) and a maximum Z-projection of Citrine-CENP-A or -CENP-C are presented. Note that Citrine-CENP-C (B) brightness/contrast were enhanced to confirm no centromeric signals at 65 min. White stars label autofluorescent chloroplasts and yellow dotted lines mark the position of the nucleus. Bars, 5 µm. (C) Relative intensity plot of Citrine signals at the centromeres and at the non-centromeric region in the nucleus (background measurement). Each line represents average relative fluorescent intensity of ≥ 6 centromeres or ≥ 6 non-centromeric regions inside the nucleus in a single cell (four cells analyzed for both Citrine-CENP-A and Citrine-CENP-C lines), measured every 15 min from the maximum Z-projection. Note that we could not identify centromeric Citrine-CENP-C signals during ∼2 h after mitotic exit, and therefore, the data are missing from the graph.

**Figure supplement 5.**
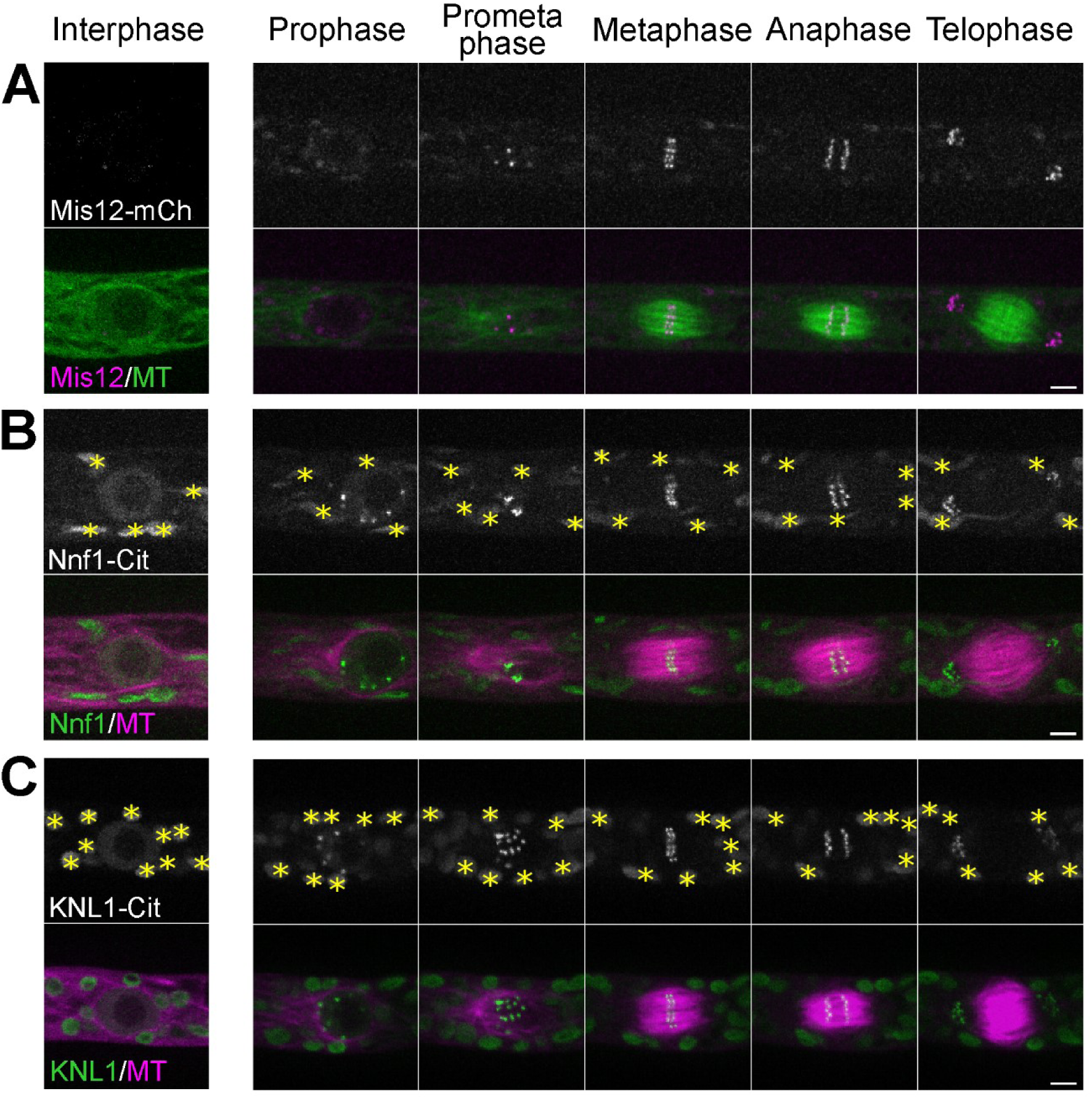
Localization of Mis12, Nnf1 and KNL1 during cell division. Live imaging of *P. patens* protonemal apical cells expressing GFP-tubulin and Mis12-mCherry (A) or mCherry-tubulin and Nnf1-Citrine(B) or KNL1-Citrine (C) KNL1-Citrine data is an expanded version of Figure 1. Autofluorescent chloroplasts are marked with yellow asterisks. Images were acquired at a single focal plane. Bars, 5 µm.

**Figure supplement 6.**
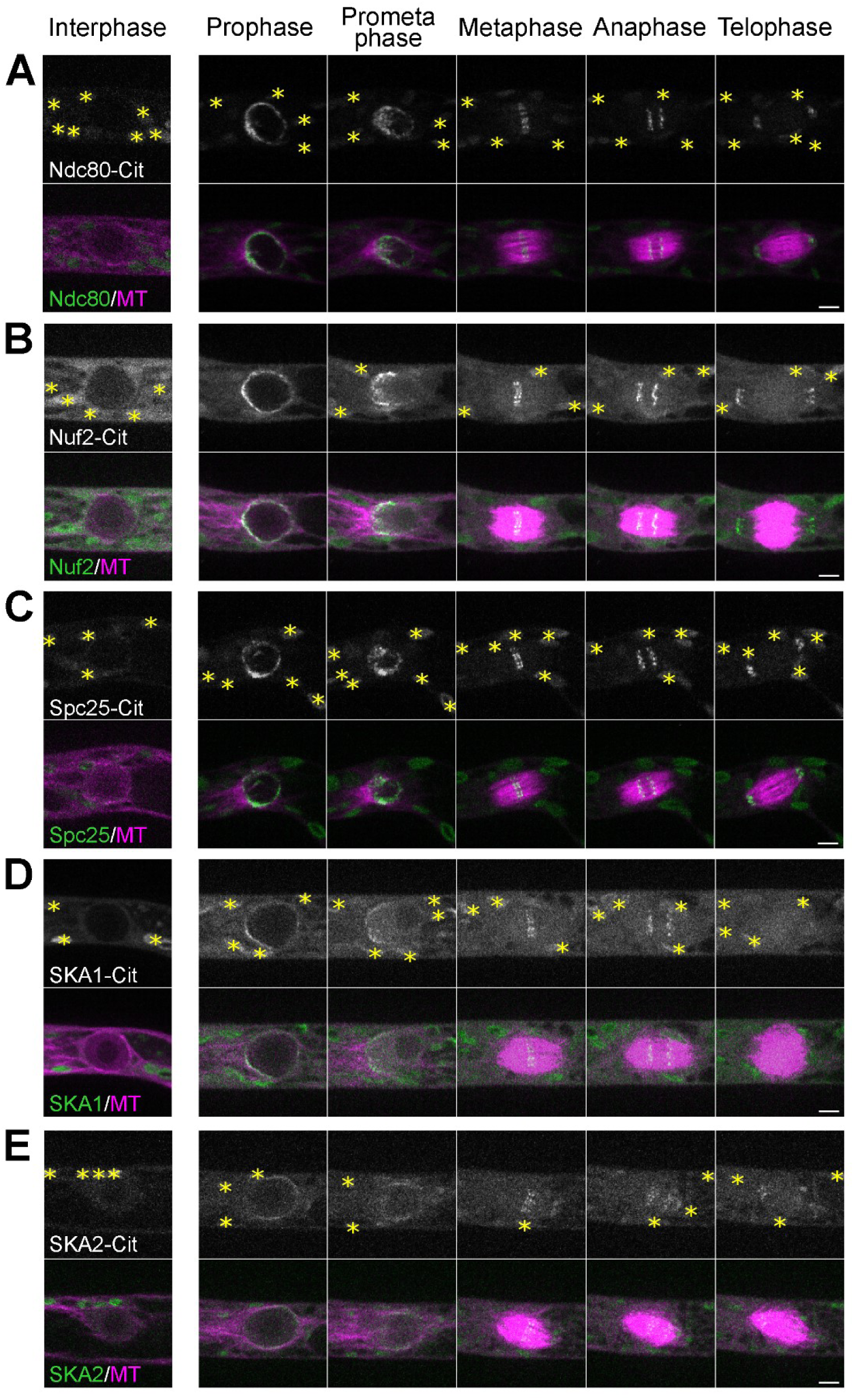
Localization of outer kinetochore proteins during cell division. Live imaging in *P. patens* protonemal apical cells expressing mCherry-tubulin (magenta) and Citrine-tagged (green) Ndc80 (A), Nuf2 (B), Spc25 (C), SKA1 (D) and SKA2 (E). Ndc80-Citrine and SKA1-Citrine data are expanded versions of Figure 1. Autofluorescent chloroplasts are marked with yellow asterisks. Images were acquired at a single focal plane. Punctate Citrine signals appeared after prometaphase. Bars, 5 µm.

**Figure supplement 7.**
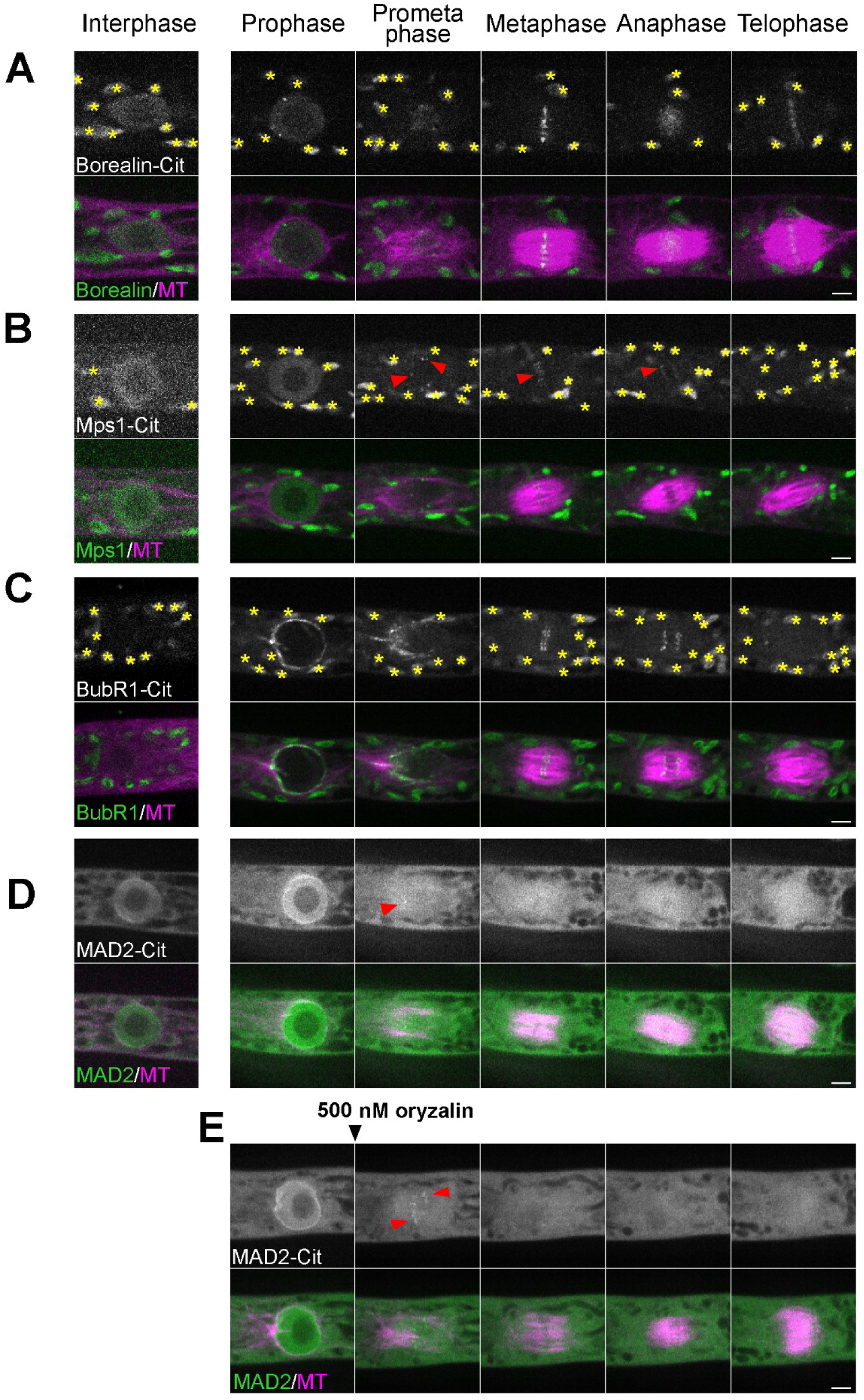
Localization of CPC and SAC proteins during cell division. Live imaging of *P. patens* protonemal apical cells expressing mCherry-tubulin (magenta) and Citrine-tagged (green) Borealin (A), Mps1 (B), BubR1(C) and Mad2 (D, E). Red arrowheads indicate punctate signals. Note that kinetochore localization of Mad2 was more clearly observed following addition of the microtubule-depolymerizing drug (500 nM oryzalin) (E). Autofluorescent chloroplasts were marked with yellow asterisks. Images were acquired at a single focal plane. Bars, 5 µm.

**Figure supplement 8.**
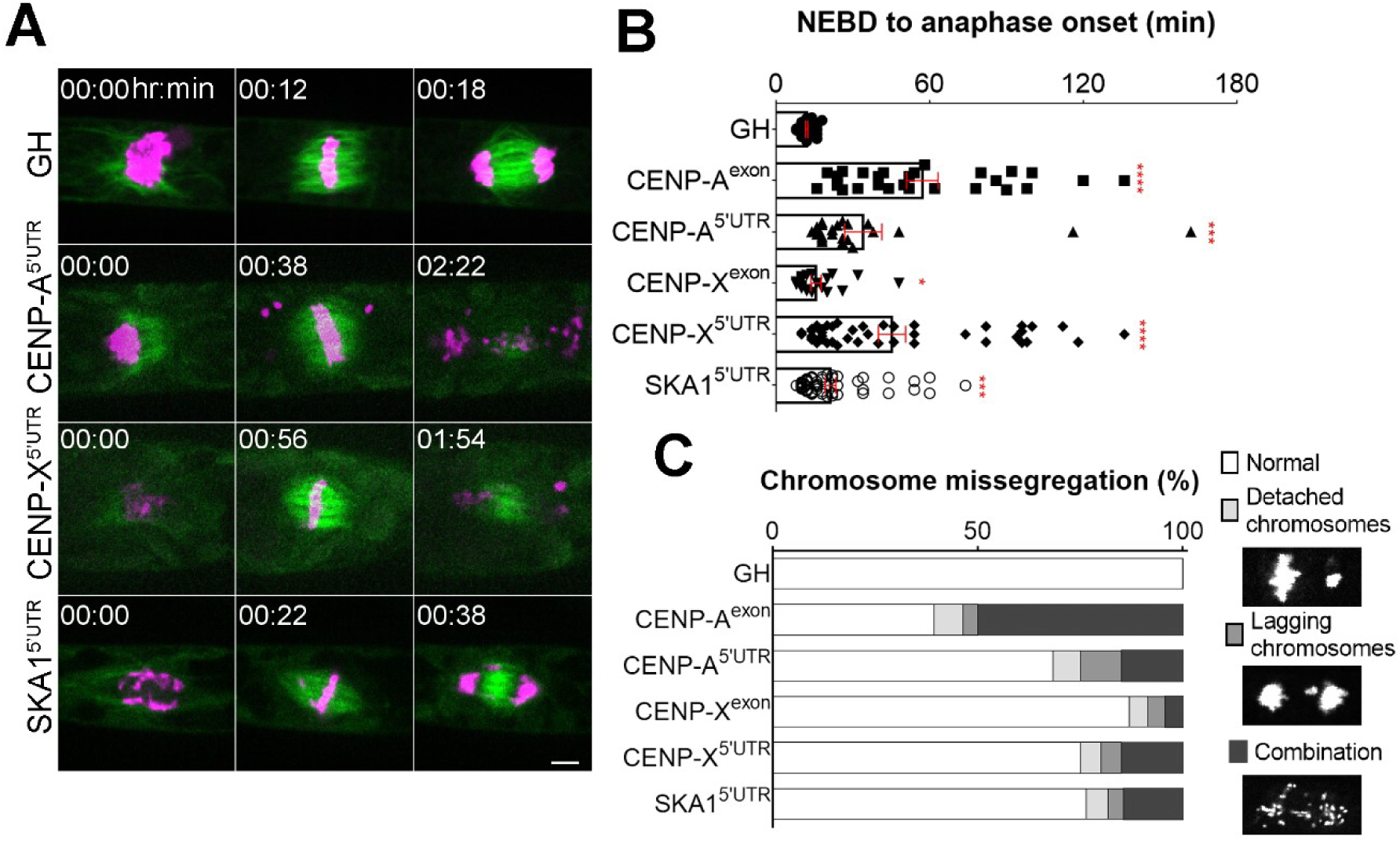
Chromosome segregation defects following depletion of CENP-A, CENP-X or SKA1. (A) Representative mitotic progression and chromosome missegregation caused by depletion of CENP-A, CENP-X or SKA1. “GH” is the control line. Bar, 5 µm. (B) Duration of mitosis (from NEBD to anaphase onset) was calculated from high-resolution live-cell imaging data for each RNAi line and the control line (GH). Bars indicate mean and SEM, whereas asterisks indicate significant differences compared with the control (*P < 0.04, ***P < 0.0007, ****P < 0.0001; two-tailed *t*-test). More than 20 cells were analyzed for each line. (C) Frequency of chromosome missegregation in different RNAi lines. Chromosome missegregation defects were classified into three types: chromosomes detached from the metaphase plate (detached chromosomes), lagging chromosomes in anaphase (lagging chromosomes), and their combination. More than 20 cells were analyzed for each line.

**Figure supplement 9.**
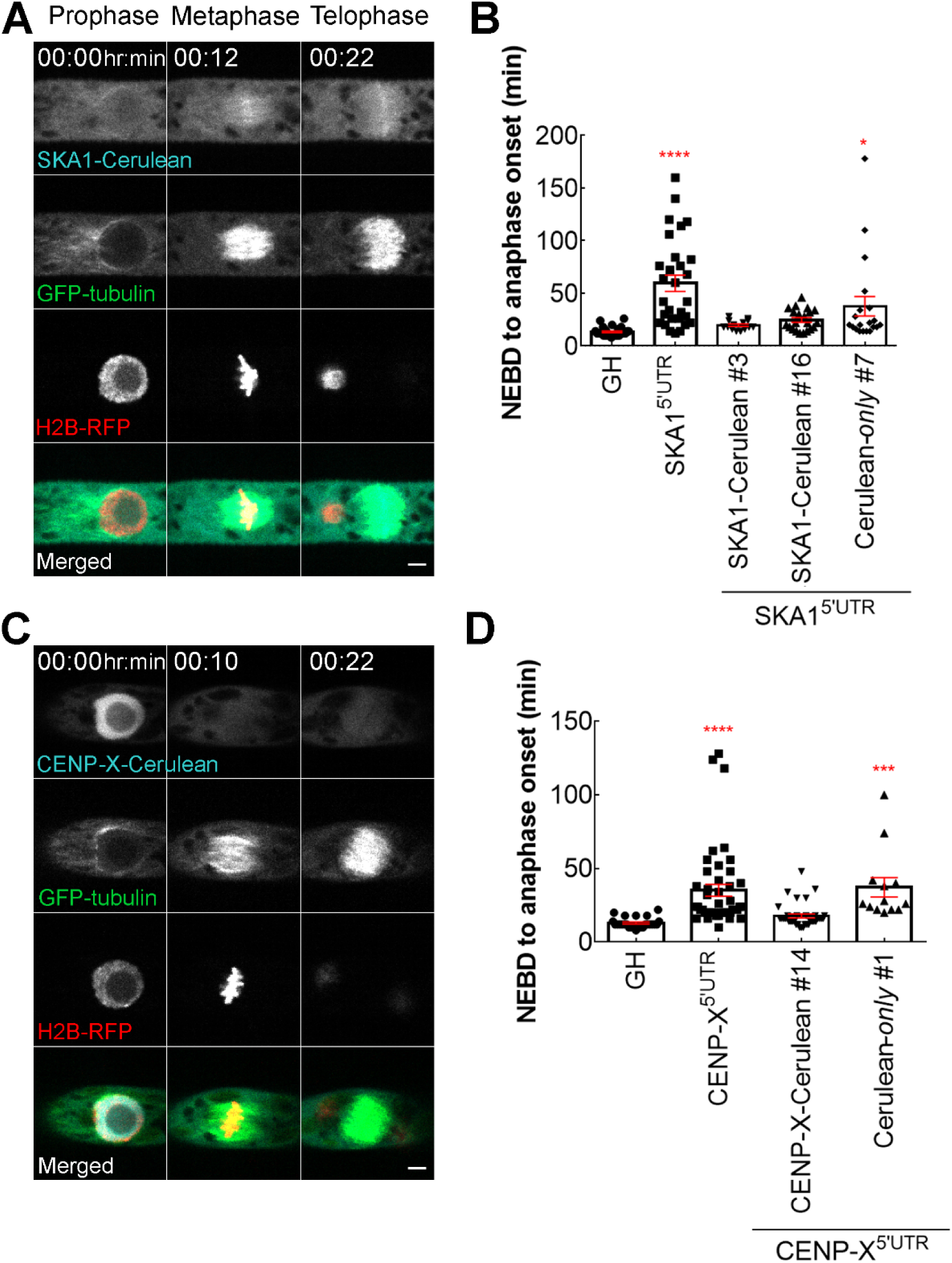
Rescue of RNAi phenotypes by ectopic expression of SKA1-Cerulean or CENP-X-Cerulean. Live imaging of *P. patens* protonemal apical cells expressing SKA1-Cerulean (A) or CENP-X-Cerulean (C) in the SKA1 5’UTR RNAi or CENP-X 5’UTR RNAi lines, respectively. RNAi was induced by addition of β-estradiol to the culture medium at the final concentration of 5 µM, 5–6 days prior to observation. Bar, 5 µm. (B, D) Mitotic duration (from NEBD to anaphase onset) for each RNAi line with or without the rescue construct (two independent SKA1 rescue lines [#3, #16] were analyzed). “GH” is the mother line used for RNAi transformation. Bars indicate mean and SEM, whereas asterisks indicate significant differences (*P < 0.03, ***P < 0.001, ****P < 0.0001; one-way ANOVA). More than ten cells were analyzed for each line.

**Table supplement 1.** Dataset used for Fisher’s test in Figure 2C.

| **CENP-A<sup>exon</sup>; CENP-A<sup>5'UTR</sup>** | Cytokinesis defect | Cytokinesis complete |
| --- | --- | --- |
| Lagging chromosomes observed in the midzone for $\leq 4$ min | 0 | 8 |
| Lagging chromosomes observed in the midzone for $\geq 12$ min | 9 | 0 |

| **CENP-X<sup>5'UTR</sup>** | Cytokinesis defect | Cytokinesis complete |
| --- | --- | --- |
| Lagging chromosomes observed in the midzone for $\leq 4$ min | 0 | 4 |
| Lagging chromosomes observed in the midzone for $\geq 8$ min | 14 | 0 |

| **SKA1<sup>5'UTR</sup>** | Cytokinesis defect | Cytokinesis complete |
| --- | --- | --- |
| Lagging chromosomes observed in the midzone for $\leq 4$ min | 0 | 3 |
| Lagging chromosomes observed in the midzone for $\geq 6$ min | 5 | 1 |

**Video 1. Localization of the centromere and CCAN proteins during cell division**

Live-cell imaging was conducted in *P. patens* protonemal cells expressing mCherry-tubulin (magenta) and one of the following tagged proteins(green): Citrine-CENP-A, KNL2-Citrine, Citrine-CENP-C, Citrine-CENP-O and Citrine-CENP-S. Note that brightness/contrast of Citrine-CENP-O images have been enhanced. Images are single focal plane and were acquired every 30 s. Bar, 10 µm.

**Video 2. Transient disappearance of CENP-C from the kinetochore after cell division**

Live-cell imaging was conducted in *P. patens* protonemal cells expressing mCherry-tubulin (magenta) and one of the following tagged proteins (green): Citrine-CENP-A, Citrine-CENP-C and KNL2-Citrine. Displayed are the the merged images of a single focal plane for mCherry-tubulin (magenta) and maximum-projection of the Z-stack for Citrine-tagged proteins. Images were acquired every 5 min. Bar, 10 µm.

**Video 3. Localization of the C-termini tagged CENP-C and CENP-O**

Live-cell imaging was conducted in *P. patens* protonemal cells expressing mCherry-tubulin (magenta) and one of the following tagged proteins(green):CENP-C-Citrine and CENP-O-Citrine. Images are single focal plane and were acquired every 30 s. Bar, 10 µm.

**Video 4. Localization of the Mis12, KNL1, Nuf2 and SKA1 during cell division**

Live-cell imaging was conducted in *P. patens* protonemal cells expressing mCherry-tubulin or GFP-tubulin (magenta) and one of the following tagged proteins: Mis12-mCherry, KNL1-Citrine, Nuf2-Citrine and SKA1-Citrine Images were acquired at a single focal plane every 30 s. Bar, 10 µm.

**Video 5. Mitotic defects observed in RNAi lines targeting CENP-C, Nnf1, Nuf2 and KNL1**

Representative images of mitotic progression and defects caused by depletion of four kinetochore proteins. White boxes indicate normal cell division in the control line (GH). White arrowheads show position of multinucleated cells, yellow arrowheads indicate chromosome missegregation and cytokinesis failure events, whereas cyan arrowheads show dead cells. RNAi was induced by addition of β-estradiol to the growth medium at the final concentration of 5 µM, 5–6 days prior to observation. Images were acquired at a single focal plane every 3 min. Bar, 100 µm.

**Video 6. Chromosome missegregation after RNAi**

Representative images of mitotic progression and chromosome missegregation caused by depletion of CENP-A or CENP-X or SKA1. RNAi was induced by addition of β-estradiol to the growth medium at final concentration of 5 µM, 5–6 days prior to observation. Images were acquired at a single focal plane every 2 min. Bar, 10 µm.

**Video 7. Cytokinesis defect associated with lagging chromosomes in anaphase**

Representative images of correlation between lagging chromosomes and cytokinesis defect in CENP-A exon RNAi and SKA1 5’UTR RNAi lines. Note that minor lagging chromosomes observed in the midzone for ≤ 4 min did not affect cytokinesis (*upper rows*); however lagging chromosomes persistent for ≥ 6 min resulted in cytokinesis failure (*bottom rows*). This correlation is conserved in both CENP-A exon RNAi and SKA1 5’UTR RNAi lines. Cytokinesis failure was concluded when the nucleus moved without restraint of the cell plate. RNAi was induced by addition of β-estradiol to the growth medium at final concentration of 5 µM, 5–6 days prior to observation. Images were acquired at a single focal plane every 2 min. Bar, 10 µm.

**Video 8. Visualization of the cell plate formation using FM4-64 dye**

Representative images of cytokinesis in the control GH line *(upper row)*, SKA1 5’UTR RNAi line with minor lagging chromosomes *(middle row),* and with persistent lagging chromosomes *(bottom row)*. Cell plate formation was visualized with 25 µM endocytic FM4-64 dye added during metaphase. FM4-64 dye was prone to photobleaching, and therefore was sometimes supplied multiple times during long-term imaging *(bottom row)*. Images were acquired at a single focal plane every 2 min. Bar, 10 µm.

**Video 9. Mitotic entry of the multi-nucleated cell in *P. patens***

SKA1 5’UTR RNAi was induced by addition of β-estradiol to the growth medium at final concentration of 5 µM, 5–6 days prior to observation. Multi-nucleated cells resulting from cytokinesis failure were monitored with the spinning-disk confocal microscope. Images were acquired at a single focal plane every 5 min. Bar, 10 µm.

**Supplemental dataset 1. *Physcomitrella patens* transgenic lines generated in this study**

**Supplemental dataset 2. Plasmids and primers used in this study**

**Supplemental dataset 3. Protein alignments used for the phylogeny analysis**

## References

1. Potapova T, Gorbsky G (2017) The Consequences of Chromosome Segregation Errors in Mitosis and Meiosis. Biology (Basel) 6(1). doi:10.3390/biology6010012.

2. Musacchio A, Desai A (2017) A Molecular View of Kinetochore Assembly and Function. Biology (Basel) 6(5). doi:10.3390/biology6010005.

3. Yu H, Hiatt EN, Dawe RK (2000) The plant kinetochore. Trends Plant Sci 5(12):543– 547.

4. Hooff JJE Van, Kops GJPL, Tromer E, Wijk LM Van (2017) Evolutionary dynamics of the kinetochore network in eukaryotes as revealed by comparative genomics. EMBO Rep 18(9):1559–1571.

5. Yamada M, Goshima G (2017) Mitotic Spindle Assembly in Land Plants: Molecules and Mechanisms. Biology (Basel) 6(1). doi:10.3390/biology6010006.

6. Jinwoo Shin, Goowon Jeong, Jong-Yoon Park HK and IL (2018) MUN (MERISTEM UNSTRUCTURED), encoding a SPC24 homolog of NDC80 kinetochore complex, affects development through cell division in Arabidopsis thaliana. Plant J 12(10):3218–3221.

7. Zhang H, et al. (2018) Role of the BUB3 protein in phragmoplast microtubule reorganization during cytokinesis. Nat Plants 4(7):485–494.

8. Wang M, et al. (2012) BRK1, a Bub1-Related Kinase, Is Essential for Generating Proper Tension between Homologous Kinetochores at Metaphase I of Rice Meiosis. Plant Cell 24(12):4961–4973.

9. Paganelli L, Lecomte P, Deslandes L (2009) Spindle Assembly Checkpoint Protein Dynamics Reveal Conserved and Unsuspected Roles in Plant Cell Division. PLoS One 4(8). doi:10.1371/journal.pone.0006757.

10. Komaki S, Schnittger A (2017) The Spindle Assembly Checkpoint in Arabidopsis Is Rapidly Shut Off during Severe Stress. Dev Cell 43(2):172–185.

11. Lermontova I, et al. (2013) Arabidopsis KINETOCHORE NULL2 Is an Upstream Component for Centromeric Histone H3 Variant cenH3 Deposition at Centromeres. Plant Cell 25(9):3389–3404.

12. Sandmann M, et al. (2017) Targeting of Arabidopsis KNL2 to Centromeres Depends on the Conserved CENPC-k Motif in Its C Terminus. Plant Cell 29(1):144–155.

13. Sato H, Shibata F, Murata M (2005) Characterization of a Mis12 homologue in Arabidopsis thaliana. Chromosom Res 13(8):827–834.

14. Du Y, Dawe RK (2007) Maize NDC80 is a constitutive feature of the central kinetochore. Chromosom Res 15(6):767–775.

15. Ogura Y, et al. (2004) Characterization of a CENP-C homolog in Arabidopsis thaliana. Genes Genet Syst 79(3):139–144.

16. Cove D, Bezanilla M, Harries P, Quatrano R (2006) Mosses as Model Systems for the Study of Metabolism and Development. Annu Rev Plant Biol 57:497–520.

17. Sato Y, et al. (2017) Cells reprogramming to stem cells inhibit the reprogramming of adjacent cells in the moss Physcomitrella patens. Sci Rep 7(1). doi:10.1038/s41598-017-01786-1.

18. Ishikawa M, et al. (2011) Physcomitrella Cyclin-Dependent Kinase A Links Cell Cycle Reactivation to Other Cellular Changes during Reprogramming of Leaf Cells C W OA. Plant Cell 23(8):2924–2938.

19. Yamada, Moe, Miki T, Goshima G (2016) Imaging Mitosis in the Moss Physcomitrella patens. Methods Mol Biol 1413:293–326.

20. Nakaoka Y, et al. (2012) An Inducible RNA Interference System in Physcomitrella patens Reveals a Dominant Role of Augmin in Phragmoplast Microtubule Generation. Plant Cell 24(4):1478–1493.

21. Miki T, Nakaoka Y, Goshima G (2016) Live Cell Microscopy-Based RNAi Screening in the Moss Physcomitrella patens. Methods Mol Biol 1470:225–246.

22. Kosetsu K, Keijzer J De, Janson ME, Goshima G (2013) MICROTUBULE-ASSOCIATED PROTEIN65 Is Essential for Maintenance of Phragmoplast Bipolarity and Formation of the Cell Plate in Physcomitrella patens. Plant Cell 25:4479–4492.

23. Miki T, Naito H, Nishina M, Goshima G (2014) Endogenous localizome identifies 43 mitotic kinesins in a plant cell. Proc Natl Acad Sci 111(11):1053–1061.

24. Naito H, Goshima G (2015) NACK Kinesin Is Required for Metaphase Chromosome Alignment and Cytokinesis in the Moss Physcomitrella Patens. Cell Struct Funct 41:31–41.

25. Hiwatashi Y, et al. (2008) Kinesins Are Indispensable for Interdigitation of Phragmoplast Microtubules in the Moss Physcomitrella patens. Plant Cell 20(11):3094–3106.

26. Ganem NJ, Pellman D (2007) Limiting the Proliferation of Polyploid Cells. Cell 131(3):437–440.

27. Uetake Y, Sluder G (2004) Cell cycle progression after cleavage failure: Mammalian somatic cells do not possess a “tetraploidy checkpoint.” J Cell Biol 165(5):609–615.

28. Panopoulos A, et al. (2014) Failure of cell cleavage induces senescence in tetraploid primary cells. Mol Biol Cell 25(20):3105–3118.

29. Schween G, Gorr G, Hohe A, Reski R (2003) Unique tissue-specific cell cycle in Physcomitrella. Plant Biol 5(1):50–58.

30. Schween G, Schulte J, Reski R (2005) Effect of Ploidy Level on Growth, Differentiation, and Morphology in Physcomitrella patens. Bryologist 108(1):27–35.

31. Hori, T. Tokuko Haraguchi, Yasushi Hiraoka HK and TF (2003) Dynamic behavior of Nuf2-Hec1 complex that localizes to the centrosome and centromere and is essential for mitotic progression in vertebrate cells. J Cell Sci 116(16):3347–3362.

32. Guse A, Carroll CW, Moree B, Fuller CJ, Straight AF (2012) In vitro centromere and kinetochore assembly on defined chromatin templates. Nature 477(7364):354–358.

33. Weir JR, et al. (2016) Insights from biochemical reconstitution into the architecture of human kinetochores. Nature 537(7619):249–253.

34. Amano M, et al. (2009) The CENP-S complex is essential for the stable assembly of outer kinetochore structure. J Cell Biol 186(2):173–182.

35. Chang DC, Xu N, Luo KQ (2003) Degradation of cyclin B is required for the onset of anaphase in mammalian cells. J Biol Chem 278(39):37865–37873.

36. Yang X, et al. (2011) The radially swollen 4 separase mutation of Arabidopsis thaliana blocks chromosome disjunction and disrupts the radial microtubule system in meiocytes. PLoS One 6(4). doi:10.1371/journal.pone.0019459.

37. Moschou PN, et al. (2013) The Caspase-Related Protease Separase (EXTRA SPINDLE POLES) Regulates Cell Polarity and Cytokinesis in Arabidopsis. Plant Cell 25(6):2171–2186.

38. Wu S, et al. (2010) A conditional mutation in Arabidopsis thaliana separase induces chromosome non-disjunction, aberrant morphogenesis and cyclin B1;1 stability. Development 137:953–961.

39. Norden C, et al. (2006) The NoCut Pathway Links Completion of Cytokinesis to Spindle Midzone Function to Prevent Chromosome Breakage. Cell 125(1):85–98.

40. Amaral N, et al. (2016) The Aurora-B-dependent NoCut checkpoint prevents damage of anaphase bridges after DNA replication stress. Nat Cell Biol 18(5):516–526.

41. de Keijzer J, Kieft H, Ketelaar T, Goshima G, Janson ME (2017) Shortening of Microtubule Overlap Regions Defines Membrane Delivery Sites during Plant Cytokinesis. Curr Biol 27(4):514–520.

42. Menéndez-Yuffá A, Fernandez-Da Silva R, Rios L, Xena de Enrech N (2000) Mitotic aberrations in coffee (Coffea arabica cv. ‘Catimor’) leaf explants and their derived embryogenic calli. Electron J Biotechnol 3(2):161–166.

43. Nichols C (1941) Spontaneous chromosome aberrations in Allium. Genetics 26(1):89– 100.

44. Kvitko O V., Muratova EN, Bazhina E V. (2011) Cytogenetics of Abies sibirica in decline fir stands of West Sayan High Mountains. Contemp Probl Ecol 4(6):641–646.

45. Smertenko A, et al. (2017) Plant Cytokinesis: Terminology for Structures and Processes. Trends Cell Biol 27(12):885–894.

46. Rensing S, et al. (2008) The Physcomitrella Genome Reveals Evolutionary Insights into the Conquest of Land by Plants. Science 319(5859):64–69.

47. Lopez-Obando M, et al. (2016) Simple and Efficient Targeting of Multiple Genes Through CRISPR-Cas9 in Physcomitrella patens. G3 Genes|Genomes|Genetics 6(11):3647–3653.

48. Vidali L, Augustine RC, Kleinman KP, Bezanilla M (2007) Profilin Is Essential for Tip Growth in the Moss Physcomitrella patens. Plant Cell 19(11):3705–3722.

